# Overcoming resolution loss due to thermal magnetic field fluctuations from phase plates in transmission electron microscopy

**DOI:** 10.1101/2023.02.12.528160

**Authors:** Jeremy J. Axelrod, Petar N. Petrov, Jessie T. Zhang, Jonathan Remis, Bart Buijsse, Robert M. Glaeser, Holger Mȕller

## Abstract

We identify thermal magnetic field fluctuations, caused by thermal electron motion (“Johnson noise”) in electrically conductive materials, as a potential resolution limit in transmission electron microscopy with a phase plate. Specifically, resolution loss can occur if the electron diffraction pattern is magnified to extend phase contrast to lower spatial frequencies, and if conductive materials are placed too close to the electron beam. While our initial implementation of a laser phase plate (LPP) was significantly affected by these factors, a redesign eliminated the problem and brought the performance close to the expected level. The resolution now appears to be limited by residual Johnson noise arising from the electron beam liner tube in the region of the LPP, together with the chromatic aberration of the relay optics. These two factors can be addressed during future development of the LPP.

## 1. Introduction

In single-particle electron cryo-microscopy (cryo-EM) [1-3] and tomography (cryo-ET) [4-6], a phase plate can provide optimum image contrast for weak-phase objects [7, 8] by selectively phase shifting the undiffracted part of the electron wave function by 90° relative to the diffracted part in the back focal plane of the objective lens. Although many designs have been investigated, it has been challenging to find a solution that is in practice superior to simply defocusing the image [9, 10].

We have demonstrated a laser phase plate (LPP) that is based on coherently manipulating the electron wave function with a continuous laser beam, which is built up to an intensity of ∼400 GW/cm^2^ by resonance in a Fabry-Perot cavity. Because no material objects are placed into the electron beam, the LPP can withstand indefinite electron exposure, and it is free of charging effects and unwanted electron scattering [11-13]. To overcome physical space limitations within the narrow gap of the objective lens, and to obtain a low cut-on frequency of ∼0.004 nm^-1^, we use relay optics for the electron beam (see Section 2 and Figure 1). The cut-on frequency, above which phase contrast becomes effective, is here defined as the point where the contrast transfer function exceeds a magnitude of 0.5. The relay optics generate a magnified version of the diffraction pattern and allows installing the LPP in a plane where more physical space is available.

**Figure 1.**
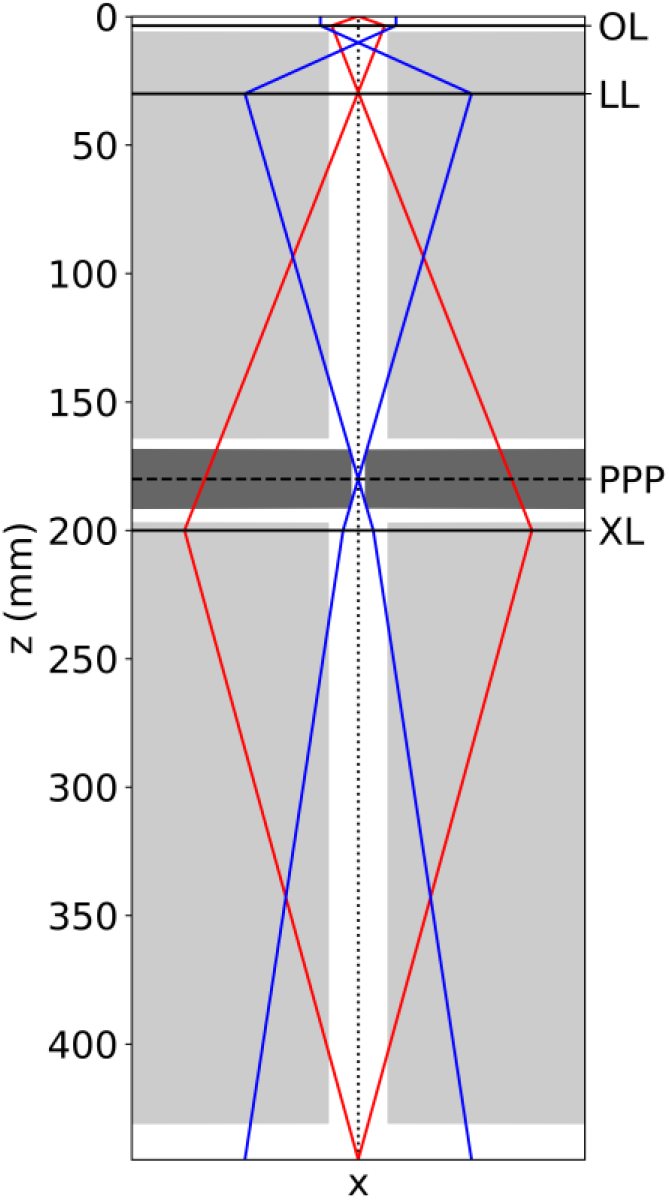
Ray diagram of the modified Titan 80-300 keV TEM used in this work. The marginal ray (red) and paraxial ray (blue) are shown between the sample plane (*z* = −0.1 mm, at the top of the Figure) and the selected area aperture plane (*z* = 445 mm, at the bottom of the Figure). The marginal ray is drawn so that it leaves the optical axis (vertical dotted line) in the sample plane at an angle of 1 radian. The Lorentz lens (LL) forms a magnified image of the back focal plane of the objective lens (OL) in the phase plate plane (PPP) at *z* = 180 mm. In that plane, the distance of the marginal ray from the optical axis, *X*(*z*), is approximately 20 mm. Cross-sections of the electron beam liner tube and LPP dummy (2 mm hole diameter) are represented by the light gray and dark gray shaded regions, respectively. The dimensions of the liner tube shown in this figure have been simplified at the request of the microscope manufacturer.

We previously were able to demonstrate the long-term stability of the LPP during data collection as well as the high contrast of images, and produced a 3-D density map of 20S proteasome particles at a resolution of 3.8 Å [14]. We initially believed that this rather modest resolution, given the number of asymmetric units included in the final reconstruction, was limited by the increased value of chromatic aberration associated with the relay optics described above. However, subsequent efforts to improve the resolution, or to reduce the number of particles needed for a given reconstruction, revealed the presence of a further limitation, in addition to the expected falloff of the temporal coherence envelope.

Here, we identify thermal magnetic field fluctuations [15, 16], which occur near the electrically conductive materials of the LPP, as the cause of the unexpected falloff of signal at high resolution. These magnetic field fluctuations are already known to potentially limit the resolution in aberration-corrected transmission electron microscopy, and though they have negligible effects in standard TEMs, their effect is enhanced here because of the magnification from the relay optics and the fact that the electron beam has to pass through a long (23 mm) and relatively narrow (2 mm diameter) hole in our initial LPP prototype. Using recombinant heavy chain human apoferritin as a test specimen, we first demonstrate a higher resolution of 2.9 Å when using the relay optics without an LPP inserted. We then present a theory-motivated redesign of the LPP, which is intended to reduce the magnitude of the magnetic field fluctuations by using a larger-diameter hole for the electron beam. Finally, we show that using mechanical (but not functional) dummy prototypes of this redesigned LPP, the resolution of the microscope is restored to a value close to that achieved without an LPP by increasing the diameter of the electron beam hole to 8 mm.

## 2. Relay-optical design of our laser phase plate microscope

The modified Thermo Fisher Scientific “Titan” 80-300 keV transmission electron microscope (TEM) used for development of the laser phase plate has been described previously [14, 17, 18]. A 5.7-fold magnified version of the electron diffraction pattern is relayed into a phase plate module, which is located below the octagon in the microscope column. When using the relay optics, the standard Lorentz lens (focal length 22 mm) is used as a first transfer lens, creating a magnified diffraction pattern 180 mm below the back focal plane of the objective lens (Figure 1). In that plane, two 25 mm diameter ports in the microscope column, at nearly opposite locations with respect to each other, allow installation of the phase plate. An additional “X-lens” located below the laser phase plate serves as a second transfer lens to pass a real-space image on to the standard projection lens optics.

The relationship *s* = *r*/*λ*_*e*_*f*_*eff*_ between spatial frequency *s* (measured in the specimen plane), electron wavelength *λ*_*e*_, effective focal length of the objective *f*_*eff*_ = 20 mm and radius *r* of features in the Fourier transform plane predicts that the cut-on spatial frequency (at which phase contrast first becomes effective) scales like the radius at which the relative phase shift first rises to an appreciable fraction of 90°. The cut-on frequency for a laser phase plate is determined by the intensity distribution of the focused light, which is not rotationally symmetric in the plane of the phase plate [18]. Regardless, azimuthally averaged values of the contrast transfer function (CTF) first exceed 50% at a spatial frequency of ∼0.004 nm^-1^, which is sufficient to image even large macromolecular complexes. Beyond this cut-on, the azimuthal average of the CTF approaches nearly 100% at higher spatial frequencies; for example, it is above 80% at 0.2 nm^-1^.

This relay optics system maintains a relatively low coefficient of spherical aberration, *C*_*s*_: the Lorentz lens contributes negligibly to *C*_*s*_, being located near a plane where the marginal ray crosses the optical axis, while the X-lens operates under low numerical aperture and so only contributes modestly. Unfortunately, however, the chromatic aberration caused by the relay optics cannot be reduced in a similar manner. All in all, the combination provides an effective focal length *f*_*eff*_ = 20 mm (compared to 3.5 mm for the objective lens alone), *C*_*s*_ = 4.7 mm (compared to 2.7 mm), and an increased chromatic aberration coefficient of *C*_*c*_ = 7.6 mm (compared to 2.7 mm). The microscope can also be operated with the relay optics deactivated, where it has the parameters of a standard Titan TEM.

## 3. Thermal magnetic field fluctuations

Uhlemann and co-authors [15, 16] have demonstrated that magnetic field noise [19-23] from electrical currents driven by thermal fluctuations in the electrically conductive parts of the instrument (Johnson-Nyquist noise, or Johnson noise for brevity) causes a loss of resolution. A magnetic field *B*_⊥_ applied over a length *L* in a direction transverse to the electron velocity will deflect the electron beam by an angle *θ* = *ηB*_⊥_*L*, where *η* = *λ*_*e*_*e*/*h, λ*_*e*_ is the electron wavelength, *e* is the elementary charge, and *h* is the Planck constant. This expression is valid for small angles *θ* ≪ 1. If the magnetic field varies stochastically in both space and time, then the variance (over time) in deflection angle ⟨*θ*^2^⟩ will depend on the magnetic field’s spatial correlation length *ξ*:

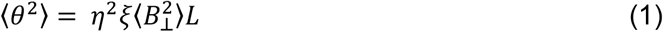

While detailed models of the spatiotemporal structure of the Johnson noise requires finite-element numerical calculations, an analytic result exists for the product *ξ*⟨*B*^2^⟩ when an electron beam passes through a tube of length *L* and diameter *D*, where *L* ≫ *D*, and the tube is made of conducting, non-magnetic material [16]:

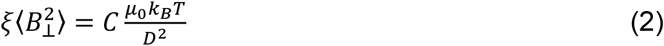

where *C* ≈ 0.77 is a dimensionless coefficient, *µ*_0_ is the permeability of free space, *k*_*B*_ is Boltzmann’s constant, and *T* is the temperature of the tube.

In our microscope, the diameter *D* of the electron beam liner tube (including phase plate, if present) changes as a function of distance *z* along the optical axis. To obtain a simple model that allows us to estimate the cumulative effect of Johnson noise along the length of the liner tube without resorting to numerical simulations, we postulate a differential formulation of Equation 1,

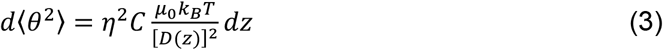

where the diameter of the tube is now a function of *z*.

The angular deflection encountered in each *z*-location is then translated into a corresponding change in apparent object position *δx* using ray transfer matrix analysis (see Section S1). The resulting variance in apparent object position due to Johnson noise then becomes an integral over the length of the electron beam liner tube:

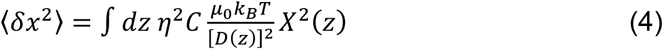

where *X*(*z*) is the distance between the electron beam’s optical axis and the marginal ray which crosses the optical axis in the object plane at an angle of 1 rad.

We assume that this variance is generated by a Gaussian distribution 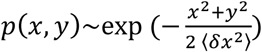, so that the image – when blurred by Johnson noise – can be written as the convolution

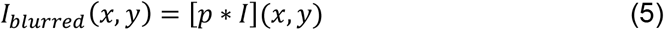

where *I* represents the unblurred image. Thus, the Fourier transform of the blurred image becomes

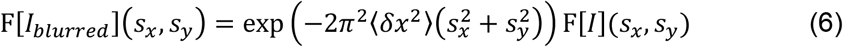

where *s*_*x*_ and *s*_*y*_ are the Cartesian components of the spatial frequency. To summarize, the blurring from Johnson noise results in the CTF being multiplied by an additional envelope function

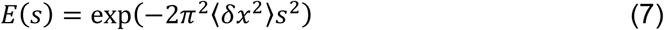

Where 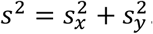.

## 4. Methods

### 4.1 Single-particle cryo-EM data collection without a laser phase plate

VitroEase™ Apoferritin Standard (Catalog number: A51362, Thermo Fisher Scientific) was used as supplied, without further purification, to prepare thin vitreous samples on Quantifoil type R 1.2/1.3 holey-carbon grids (Quantifoil Microtools). The grids were made hydrophilic, immediately before the sample was applied, by exposure to a glow discharge (PELCO easiGlow). A Mark IV Vitrobot (FEI), pre-equilibrated at 4 °C and 100 % relative humidity, was used to vitrify samples. Grids were blotted for 4 s with a blot force of 15 and then plunged immediately into liquid ethane, without pausing after blotting.

Images of vitrified specimens were recorded at 300 keV both with and without using the relay optics, using conditions that otherwise were as identical as possible. A Gatan 626 cryo-transfer holder was used to hold the sample in the TEM, and a Gatan K2 camera was used to record images as a series of movie frames. The pixel size was chosen to be close to 0.5 Å (referred to the specimen) in order to improve the detective quantum efficiency (DQE) at the band limit of the specimen. A total of 50 frames were aligned and summed to produce final values of ∼50 e/Å^2^ for the accumulated electron exposures.

Although our Titan is fitted with a cryo-box, we found a high ice accumulation rate on the specimens during imaging. Therefore, a 200 µm diameter objective lens aperture was inserted and further vacuum pumping capacity was also added to the column in the region of the X-lens. These changes reduced the observed ice buildup rate by nearly an order of magnitude. Further documentation about the improvement in contamination rate achieved by using an objective aperture is provided in Section S2.

Serial EM [24] was used to manage data acquisition. The diameter of the illuminated area was set to be ∼0.4 µm larger than the hole size in the Quantifoil grids, to minimize the effect of specimen charging. This meant, of course, that we were not able to benefit from using beam-image shift to record more than a single image per hole. Even so, approximately 10,000 candidate particles were boxed from approximately 110 movies that were collected over a period of approximately 3 hours.

### 4.2 Three-dimensional reconstruction of cryo-EM density maps

The RELION software package [25] was used for data processing and three-dimensional reconstruction. Although a larger number of candidate particles was initially extracted for each data set, 3D classification subsequently reduced this to approximately 6000 particles (144,000 asymmetric units). Only these particles were included within the final data sets used to produce the refined density maps.

### 4.3 Characterizing the CTF envelope function when using different electron beam hole diameters in the phase plate dummies

To characterize the effect of different hole diameters, simplified mechanical dummies were inserted into the port on the microscope’s phase plate module (described in Section 2). These dummies consisted of a solid aluminum cylinder (indicated by the dark gray shaded region in Figure 1) with a diameter identical to that of our actual LPP (23 mm). Each dummy had a single hole through which the electron beam could pass. The axis of that hole was perpendicular to and intersected with the axis of the dummy’s cylinder similar to the actual LPP (and such that the length of the hole was equal to the diameter of the holder). Dummies with electron beam hole diameters of 2 mm, 4 mm, and 8 mm were used. The test was also performed with no dummy inserted.

2 nm thick carbon films supported on Quantifoil holey carbon grids (Ted Pella 668-300-Cu) were imaged with a room-temperature specimen holder in order to make quantitative measurements of the microscope’s CTF envelope as a function of the diameter of the electron beam hole in the dummies. Images were recorded as movies consisting of 150 frames each, using a Gatan K2 camera, with a pixel size (referred to the specimen) of 0.28 Å with relay optics and 0.32 Å without relay optics. Approximately 50 such images were collected in both modes for each size of hole. The accumulated electron exposure at the sample was ∼2300 e/Å^2^ per movie, while the dose rate at the camera was maintained at ∼8 e/pix/s in order to ensure minimal coincidence loss [26].

The CTF envelope with relay optics was estimated by computing the ratio of the background-subtracted power spectra of images taken with relay optics to those taken without relay optics. Further details of how envelopes were fitted to the power spectra are given in Section S3. This ratio was used to factor out the unwanted contributions of the camera DQE and the structure factor of the specimen, which are expected to be the same in the two imaging modes. Estimates of the envelope with relay optics were then obtained by multiplying by the theoretical values of the temporal coherence envelope without relay optics. The estimated CTF envelope with relay optics is expected to reflect a fixed contribution due to the increased amount of chromatic aberration (discussed in Section 2) and a hole diameter-dependent contribution due to Johnson noise.

### 4.4 Estimating the image blur that corresponds to an envelope function for a given hole diameter

According to Equation 7, Johnson noise generates a Gaussian contribution to the falloff of the CTF envelope. We also take into account chromatic aberration by including the usual temporal coherence envelope (see Section S4), the exponent of which increases as the fourth power of the spatial frequency. The product of these two envelopes was used to fit the measured CTF envelope data, using the image blur variance ⟨*δx*^2^⟩ and electron beam energy spread as fit parameters. The electron beam energy spread fit parameter was forced to be the same for all four measured CTF envelopes (i.e. for the three electron beam hole diameters and the case where no dummy was inserted). To summarize, five parameters were used to fit the four measured CTF envelopes—four representing the different image blur variances, and one representing the identical temporal coherence envelopes.

## 5. Results

### 5.1 High quality cryo-EM results are achieved when a laser phase plate is not inserted

Cryo-EM images of apoferritin particles were collected both with and without relay optics (without a phase plate inserted in either case), to establish a baseline of the performance that might be achieved with our phase plate development microscope. Examples of the images that were obtained with defocus values of approximately 1.2 µm are shown in Figure S2, while examples of fits of the known structure of apoferritin (PDB accession number 6Z6U), docked as a rigid body into the refined density maps, are shown in Figure 2A,B.

**Figure 2.**
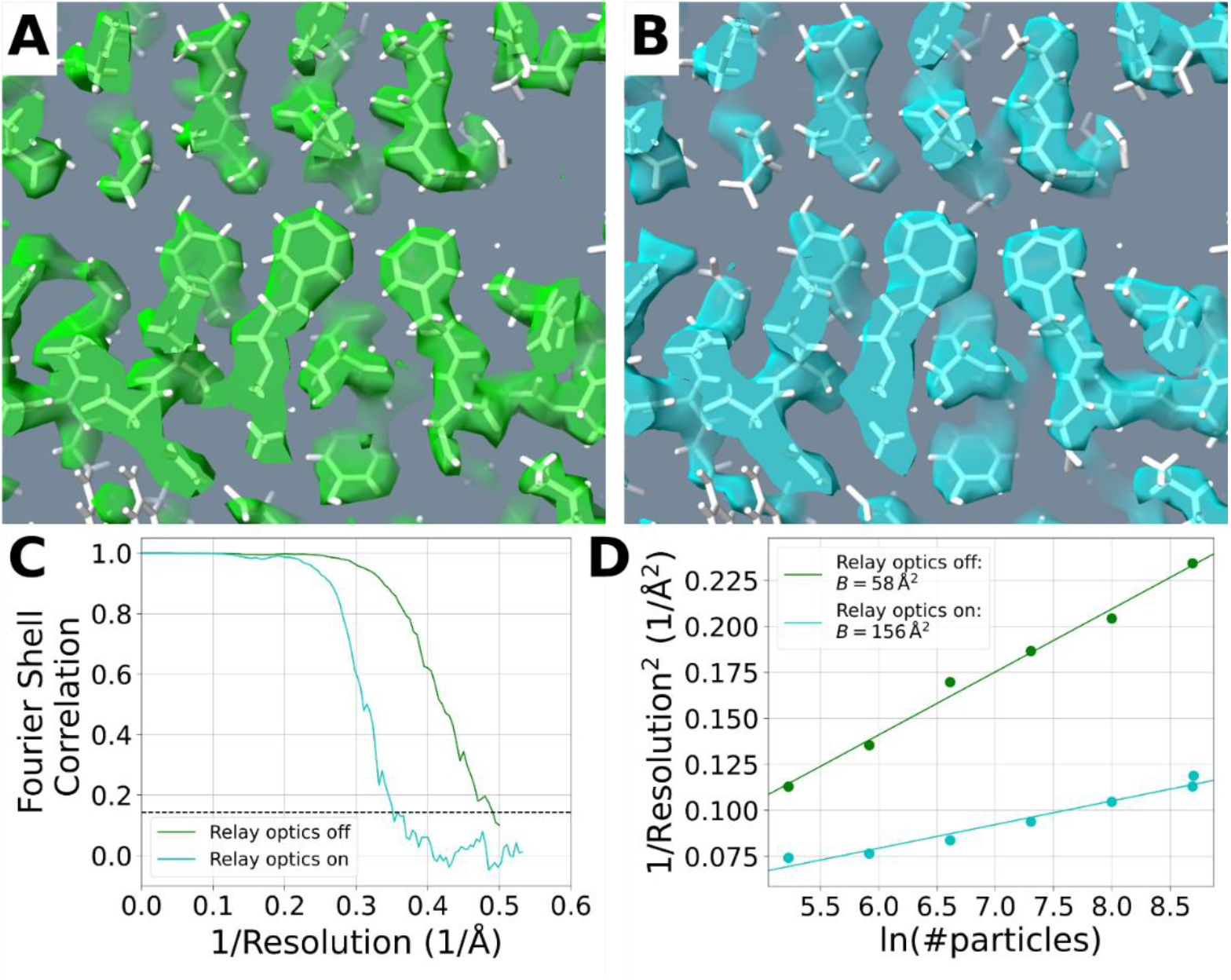
Benchmarking the relative performance of the modified Titan that is currently used for development of a laser phase plate. **(A)** Example of a portion of the refined density map of apoferritin, which was obtained without relay optics and is displayed with graphics provided in Chimera [30]. The atomic model that was docked as a rigid body into the EM density was obtained from PDB accession number 6Z6U. **(B)** The same portion of the map as is shown in (A), but this time obtained with relay optics. **(C)** Comparison of the gold-standard Fourier shell correlation (FSC) curves for data collected without relay optics (green), and with relay optics (blue). The dashed line indicates the FSC = 0.143 resolution cutoff level. **(D)** Plots of the inverse-squared resolution achieved as a function of the natural logarithm of the number of particles (multiply by 24 to get the number of asymmetric units) in the data set, generated by *bfactor_plot*.*py* in Relion 3.1.3 [25]; the upper (green) line is fitted to the results obtained without relay optics, and the lower (blue) line is fitted to the results obtained with relay optics. The corresponding B-factor (two divided by the slope of the best fit line) for each dataset is listed in the legend.

The reconstruction obtained with data collected without relay optics proved to be comparable to that in the literature, when using high quality apoferritin samples and state-of-the-art electron microscopes. As is shown in Figure 2C,D, a global (gold standard [27]) resolution of ∼2.0 Å was achieved without relay optics, using 5952 particles (1428,848 asymmetric units), and a B-factor equal to ∼58 Å^2^ was obtained from the Rosenthal-Henderson plot for these data. This value of B-factor is significantly better than what was previously reported for data collected with a side-entry cryo-holder and 200 keV electron beam [28], but not as good as the value of 32 Å^2^ reported for data collected with a Krios microscope equipped with a cold field emission gun (FEG) and an imaging filter [29].

As expected, the reconstruction obtained with relay optics was affected by the increased chromatic aberration, and possibly other factors such as Johnson noise generated in the beam liner tube, which are enhanced by the longer effective focal length. As is shown in Figure 2C,D, a global (gold standard) resolution of ∼2.9 Å was achieved, this time using 6003 particles (144,072 asymmetric units), and the value of the B-factor increased to ∼156 Å^2^ for these data. Since this increase in B-factor (and the corresponding loss in resolution achieved with a fixed number of particles) can be at least partly attributed to the increased chromatic aberration inherent to the use of relay optics, we expect that the performance with relay optics could be improved by using an electron gun with a narrower energy spread; see the Discussion (Section 6).

### 5.2 The CTF envelope deteriorates when the diameter of the hole in the laser phase plate is too small

In order to determine whether Johnson noise was a factor that previously limited our first cryo-EM results, reported in [14], we measured the CTF envelopes with LPP dummies with different hole diameters, and when no dummy was inserted into the column. The results, presented in Figure 3A, confirmed that the CTF envelope is sensitive to the diameter of the hole. When a dummy with a 2 mm hole is inserted, for example, the CTF envelope at resolutions close to ∼3 Å falls almost twofold compared to when no dummy is inserted. Increasing the diameter of the hole to 8 mm restores the CTF envelope to nearly that obtained without a dummy.

**Figure 3.**
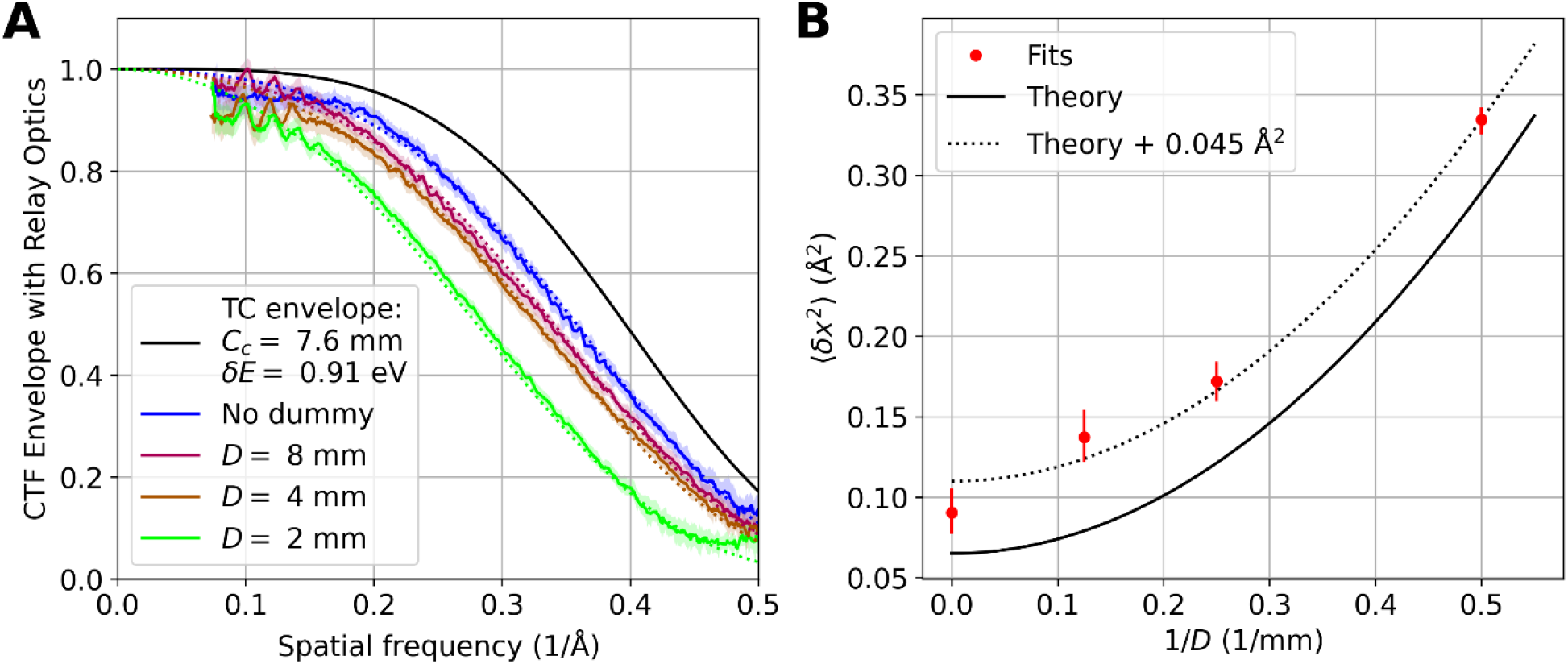
**(A)** Contrast transfer function (CTF) envelope functions with relay optics measured either without inserting a laser phase plate dummy or when dummies were inserted that had electron beam hole diameters equal to 8 mm, 4 mm, or 2 mm. The colored, solid lines represent the measured CTF envelope, with the associated shaded regions representing a 50% confidence interval. The dashed lines represent a best fit to the data, modelled as the product of the envelopes due to imperfect temporal coherence and Johnson noise. The best fit electron beam energy spread *δE* (full width half maximum) was 0.91 eV (50% confidence interval [0.88, 0.94] eV), somewhat higher than the manufacturer’s specification of 0.8 eV for the S-FEG field emission gun used here. The black solid line shows the theoretical temporal coherence (TC) CTF envelope function in the absence of Johnson noise. **(B)** The fitted values of the Johnson noise image blur variance (red dots) plotted as a function of the inverse of the dummy’s electron beam hole diameter. The error bars show the 50% confidence interval. The solid black line shows the theoretical value of the image blur variance as a function of inverse hole diameter (based on Equation 4, using manufacturer-supplied values for the marginal ray distance and liner tube diameter). The dotted black line represents a best fit to the data when a constant value is added to the theoretical model as a fit parameter (see Section 5.2).

Johnson noise in the microscope’s liner tube becomes dominant for dummy hole diameters of 8 mm and above. This is visible in the CTF envelope measured with no dummy inserted, as it falls below 0.5 at 0.35 Å^-1^, compared to 0.40 Å^-1^ for the theoretical temporal coherence envelope. The measured CTF envelopes shown in Figure 3A are well-fitted by theoretical curves (dashed lines) which includes Johnson noise as well as the increased chromatic aberration of the relay optics (see Section 4.4 for details).

Figure 3B shows the amount of Johnson noise image blur variance ⟨*δx*^2^⟩ extracted from the data in Figure 3A compared to the blur expected from the theoretical model in Equation 4. The non-zero predicted value that remains, when there is no dummy, arises from Johnson noise in the liner tube. The model reproduces the observed increase in image blur with decreasing electron beam hole size. However, the measured variance exceeds the prediction by a (roughly constant) 0.045 Å^2^. This could be due to an inaccuracy in the simplified model, or additional resolution loss that affects the microscope more strongly when using relay optics than when not (e.g. vibrations of the microscope column, or magnetic field noise from external sources).

## 6. Discussion

Thermal fluctuations in the magnetic field adjacent to any electrically conductive component, arising from Johnson-Nyquist noise, are known to be a limiting factor at resolutions beyond ∼0.5 Å [15] but are usually negligible at resolution values achieved in cryo-EM. However, they need to be addressed when relay optics are used to magnify the electron diffraction pattern at the plane of the phase plate. In particular, they become a limitation when a long, narrow hole (dimensions *L* = 23 mm and *D* = 2 mm) is provided for the electron beam to pass though the LPP, as was the case in [14].

Fortunately, theoretical modeling indicated that a hole diameter of 8 mm would largely resolve the problem. Our experimental results (Figure 3A), obtained when “dummy” LPPs (i.e. ones with geometries approximating that of the actual LPP but which did not include a laser cavity) were inserted, confirmed that the CTF envelope function becomes almost identical to that without any LPP when the diameter of the electron beam hole is increased to 8 mm. Even then, Johnson noise originating in the liner tube remains a limiting factor. In addition, the CTF envelope of our development microscope is limited by the increased chromatic aberration that is associated with the use of relay optics, as is described in Section 2. The CTF envelope resulting from the combination of these two factors is shown in Figure S3, where it is compared with that obtained when relay optics are not used (i.e. in a standard Thermo Fisher Scientific Titan or Krios).

These two limitations have been addressed in a design for a next-generation phase plate microscope. Residual Johnson noise has been minimized by increasing the diameter of the beam liner tube and by making it possible to adjust the effective focal length (and thus the marginal ray distance *X*(*z*)) of the relay optics between 12-20 mm. This reduces the influence of Johnson noise proportionally. The resulting increase in cut-on frequency will be negligible when imaging structure sizes smaller than about 50 nm and can be avoided by reducing the LPP’s laser focus waist and/or wavelength. Cooling the liner tube (e.g. with liquid nitrogen to ∼77 K) could be another way to substantially reduce the amount of resolution loss

The effect of chromatic aberration can be improved by reducing the width of the electron energy distribution with a gun monochromator or a cold FEG; at a width of 0.3 eV, the temporal coherence envelope with relay optics would become very similar to that achieved without relay optics and without a monochromator.

## ACKNOWLEDGEMENT

We thank A. Gonzalez, T. Gutierrez, and G. W. Long for making contributions to this work. This project was supported by the U.S. National Institutes of Health (Grant No. 5 R01 GM126011), Chan Zuckerberg Initiative (award number 2021-234606), Gordon and Betty Moore Foundation (Grant No. 9366), and a cooperative research and development agreement (CRADA) with Thermo Fisher Scientific (award number AWD00004352). Grant No. 5 R01 GM126011 and award number AWD00004352 were administered at Lawrence Berkeley National Laboratory under Contract No. DE-AC02-05CH11231. P.N.P. acknowledges support from a postdoctoral fellowship from the National Institute of General Medical Sciences of the National Institutes of Health under Award Number F32GM149186. The content is solely the responsibility of the authors and does not necessarily represent the official views of the National Institutes of Health.

