## Supplemental Material for "Overcoming resolution loss due to thermal magnetic field fluctuations from phase plates in transmission electron microscopy"

### S1. RAY TRANSFER ANALYSIS OF DEFLECTIONS IN A PLANE RELATIVE TO APPARENT OBJECT POSITION

The angular deflection applied to the electron beam by the Johnson noise magnetic field causes an apparent shift in the position of objects in the sample plane. The amount of shift depends not only on the amount of angular deflection, but also on where in the imaging system that deflection is applied. Here we quantify this using ray transfer matrix analysis.

First, consider a ray which emanates from the specimen ($z = 0$) at a distance and angle relative to the optical axis of $x_0$, and $\theta_0$, respectively. Represent this ray by a column vector $(x_0, \theta_0)^{\mathrm{T}}$. Further down the optical axis, the optical system transforms this ray into $U(z)(x_0, \theta_0)^{\mathrm{T}}$, where $U(z)$ is the ray transfer matrix representing the optical system between $z = 0$ and $z$. At this point, we add an angular deflection $\theta_1$, transforming the ray into $U(z)(x_0, \theta_0)^{\mathrm{T}} + (0, \theta_1)^{\mathrm{T}}$. In order to determine the corresponding apparent change in object position, we apply the inverse ray transfer matrix to determine the column vector $(x_0', \theta_0')^{\mathrm{T}}$ that would have produced the same ray in absence of magnetic field fluctuations, $(x_0', \theta_0')^{\mathrm{T}} = U^{-1}(z)\left(U(z)(x_0, \theta_0)^{\mathrm{T}} + (0, \theta_1)^{\mathrm{T}}\right)$. This shows that $\delta x_0 = x_0' - x_0$ can be calculated as $(1,0)U^{-1}(z)(0, \theta_1)^{\mathrm{T}} = \theta_1(1,0)U^{-1}(z)(0,1)^{\mathrm{T}}$. To simplify this expression further, note that a ray transfer matrix $U = \begin{pmatrix} A & B \\ C & D \end{pmatrix}$ between two planes with equal index of refraction has a determinant of unity, and thus its inverse is simply $U = \begin{pmatrix} D & -B \\ -C & A \end{pmatrix}$. Thus, $(1,0)U^{-1}(z)(0,1)^{\mathrm{T}} = -(1,0)U(z)(0,1)^{\mathrm{T}} \stackrel{\text{def}}{=} -X(z)$ is the distance of the marginal ray from the optical axis, where the marginal ray is defined to be that which leaves the optical axis in the sample plane at an angle of 1 rad. The marginal ray (when using relay optics) in our Titan is plotted schematically in Figure 1. We obtain the simple result that

$$\delta x(z) = -X(z)\theta_1(z)$$

We can then use this equation to sum in quadrature the apparent shifts in object position due to angular deflections caused by Johnson noise throughout the microscope column (Equation 4).

### S2. AN OBJECTIVE APERTURE HELPS TO REDUCE ICE CONTAMINATION ONTO SPECIMENS

Our Titan electron microscope is fitted with a cryo-pump (referred to as the “cryo-box”) that surrounds the specimen, similar to that provided in the Krios product line. Nevertheless, the ice contamination rate in our Titan is at least 30 times as high as it is in Krios microscopes (which typically achieve contamination rates of less than 0.5 Å/hr). As a result, cryo-EM specimens can be used in our Titan for only a relatively small number of hours, after which the ice contamination becomes too thick for high-resolution data collection.

Fortunately, we found that the contamination rate is decreased by roughly an order of magnitude by combining the use of an objective lens aperture together with adding more vacuum pumping capacity to the lower portion of the column of the microscope near the X-lens. The contamination rate was estimated by measuring the decrease in transmittance of as-supplied holey-carbon Quantifoil grids (mounted in the Gatan 626 cryo-holder) as a function of time. Transmittance values were measured by recording low-magnification images, in which the field of view of the K2 camera corresponded to ~30 µm at the specimen. Under the imaging conditions used, the initial transmittance of the carbon film was typically ~88%, measured as the number of electrons per unit area passing through the carbon film relative to the number per unit area passing through a hole. Figure S1 shows the changes in percent transmittance values when measured as a function of time, showing the resulting reduction in contamination rate that was achieved by the use of an objective aperture, together with the additional improvement achieved by increasing the pumping capacity.

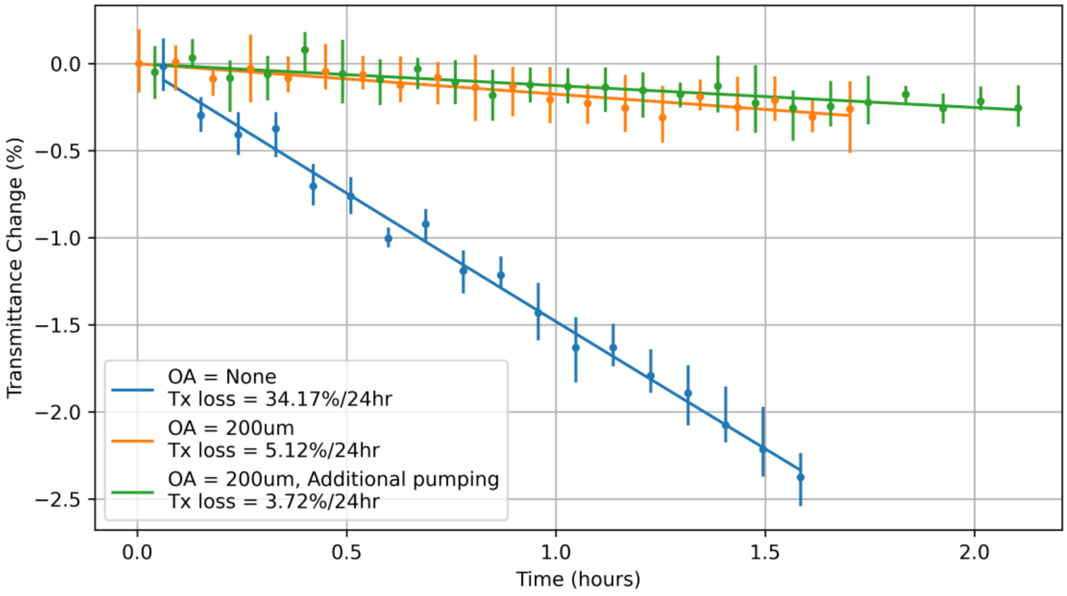


**Figure S1.** Change in percent transmittance as a function of time when i) no objective aperture (OA) is inserted and no additional vacuum pumping is used (blue), ii) a 200 µm diameter objective aperture is inserted (orange), and iii) 200 µm diameter objective aperture is inserted and additional vacuum pumping is used (green). The dots with error bars represent measurements taken at discrete intervals, while the lines represent exponential best fits to the data. The corresponding percent transmittance change rates are shown in the insert.

The biggest benefit is gained by inserting a 200 µm diameter objective lens aperture, which is inserted into the space between the cryo-box and the lower pole piece, and thus remains at room temperature. Such an aperture is large enough that inevitable charging of the aperture edge, which occurs with use, does not have a detectable effect on the Thon rings. We infer that this aperture reduces the contamination rate by reducing the solid angle over which water molecules can approach the sample from the room-temperature surfaces of the lower part of the microscope column. Though the specimen is still exposed to the surface of the room-temperature objective aperture, we expect that water molecules on this surface are quickly pumped away by the surrounding cryo-box, after which the upper, specimen-facing surface of the objective aperture, although at room temperature, is no longer a source of water molecules.

### S3. METHOD USED TO ESTIMATE THE CTF ENVELOPE FUNCTION WITH RELAY OPTICS

As described in Section 4.3 of the paper, estimation of the CTF envelope when using relay optics, relative to that when not using relay optics, was based on first computing the ratio of the background-subtracted power spectra of the respective images. Estimates of the envelope with relay optics were then obtained by multiplying this ratio by the theoretical values of the temporal coherence envelope without relay optics. This method factors out unwanted contributions of the camera DQE and the structure factor of the specimen. The resulting estimate of the CTF envelope thus reflects both a hole diameter-independent contribution due to the increased amount of chromatic aberration when using relay optics along with a hole diameter-dependent contribution, and any other factors that make the CTF envelope different with and without relay optics, if they exist.

The data collection procedure is as follows:

1. 2 nm thick carbon films supported on Quantifoil holey carbon grids (Ted Pella 668-300-Cu) were imaged with a room-temperature specimen holder.
2. The sample-referred pixel size of the images (0.28 Å with relay optics and 0.32 Å without relay optics) was calibrated using a room-temperature nanocrystalline gold sample.
3. Images were taken within the same grid square of the sample both with and without relay optics in order to reduce any systematic bias in the sample structure factor between images taken in each mode.
4. The grid hole locations of the images taken in each mode were made to form an interleaved grid pattern. This was intended to further reduce any systematic bias in the sample structure factor between images taken with and without relay optics.
5. Each grid hole was imaged only once. We found that successive images of the same area resulted in an increase in the sample structure factor (which could be produced by the build-up of hydrocarbon contamination with prolonged exposure).
6. Images were autofocused to approximately 1 µm of defocus using a region of the sample approximately 1 µm away from the target image area. This value of defocus is small enough that the associated spatial coherence CTF envelope is negligible, but large enough to generate a sufficient number of Thon rings for our data analysis procedure.
7. Images were recorded as 150 frame movies using a Gatan K2 direct electron detection camera, with a dose rate at the camera of approximately 8 e/pix/s in order to ensure minimal coincidence loss.

The data analysis procedure for each dataset (different dummy configurations) is as follows:

1. Movies from both modes were truncated (trailing frames removed) in order that all movies from the dataset had as similar as possible of a total dose per sample area. This

was done because we found that higher sample dose images resulted in higher sample structure factors.

2. The truncated movies were motion-corrected into a single image using MotionCor2 [1].
3. The CTF of the resulting motion-corrected images was then fit using CTFFIND4 [2]. In order to optimize the quality of the fit, these fits were performed as a function of the spherical aberration coefficient $C_s$ as well. This was done by running CTFFIND4 for multiple values of $C_s$ and then choosing the value which resulted in the highest 'spacing (in Angstroms) up to which CTF rings were fit successfully' output metric from CTFFIND4. The value of $C_s$ chosen this way was constrained to be the same for all subsequent image analysis within a given imaging mode (standard or phase plate).

Then, for each image in the dataset:

4. The magnitude squared of its FFT was calculated and divided by its value at zero frequency (for normalization against any remaining differences in dose between images).
5. From the normalized $|\mathrm{FFT}|^2$, the best CTF fit was used to calculate an equiphase average [3], resulting in a one-dimensional curve as a function of the mean magnitude of spatial frequency around the equiphase averaging contour.
6. The values of this curve at the spatial frequencies of the CTF zero-crossings (as determined by the CTFFIND4 best fit) were used to interpolate a two-dimensional, azimuthally symmetric function representing the frequency-dependent noise spectrum.
7. This noise floor function was then subtracted from the normalized $|\mathrm{FFT}|^2$, and the result was divided by the square of the noise-free, two-dimensional CTFFIND4 best fit function. The result should represent the power spectral density of the sample (square of the structure factor) multiplied by the square of the CTF envelope, since the expectation value of the noise spectrum has been subtracted and the oscillatory part of the CTF has been deconvolved.
8. This two-dimensional function was then equiphase averaged based on the CTFFIND4 best fit. Areas where the magnitude of the oscillatory part of the CTF was less than 0.5 were ignored as they were more sensitive to inaccuracies in the estimate of the noise floor. The resulting one-dimensional curves were linearly interpolated across these regions, and represent the envelope of the noise-subtracted power spectral density in each image.

Then, for each dataset:

9. The set of roughly 50 power spectral density curves from images taken with relay optics were divided by each of the roughly 50 power spectral density curves from images taken without relay optics, and the square root was taken. This resulted in approximately 2500 curves representing the measured ratio of the CTF envelopes, for each possible combination of images taken with and without relay optics in the dataset.

Then, for all datasets:

10. Fits (as described in Section 4.4) were performed on 10,000 random combinations of curves from each of the four datasets. They were then multiplied by the theoretical envelope without relay optics using the resulting best fit value for the electron beam energy spread $\delta E$ to generate curves representing the measured CTF envelope with relay optics.
11. The mean of these curves along with confidence interval is displayed in Figure 3A.
12. The mean and confidence interval of the image blur variance fit parameter $\langle \delta x^2 \rangle$ is shown in Figure 3B.

### S4. TEMPORAL COHERENCE CTF ENVELOPE

The temporal coherence CTF envelope is mainly due to the fact that electrons with different kinetic energies form images with different amounts of defocus, i.e. due to chromatic aberration in the microscope. The envelope function, for Gaussian distributions of the relevant parameters, is

$$\exp\left(-\frac{1}{2}\sigma_f^2\pi^2\lambda_e^2 s^4\right)$$

where $\lambda_e$ is the electron wavelength, $s$ is the spatial frequency, and

$$\sigma_f = C_c\sqrt{\left(\epsilon\frac{\sigma_{V_0}}{V_0}\right)^2 + \left(\epsilon\frac{\sigma_{E_0}}{E_0}\right)^2 + \left(2\frac{\sigma_{I_0}}{I_0}\right)^2}$$

is the spread in defocus values, where $C_c$ is the coefficient of chromatic aberration, $\epsilon = (1 + E_0/M_e)/(1 + E_0/2M_e)$ is a relativistic factor, $E_0$ is the electron's kinetic energy, $M_e$ is the rest energy of an electron, and $\frac{\sigma_{V_0}}{V_0}$, $\frac{\sigma_{E_0}}{E_0}$, and $\frac{\sigma_{I_0}}{I_0}$ are the relative standard deviations of the accelerating voltage, electron's kinetic energy when emitted from the field-emission tip, and microscope objective lens current, respectively. For this work, we used the values $\frac{\sigma_{V_0}}{V_0} = 0.07 \times 10^{-6}$ and $\frac{\sigma_{I_0}}{I_0} = 0.1 \times 10^{-6}$ though they are relatively small compared to that of $\frac{\sigma_{E_0}}{E_0} = \frac{1}{2\sqrt{2\ln 2}}\frac{\delta E}{E_0} \sim 1 \times 10^{-6}$. Recall that $\delta E$ is the full-width half-maximum of the electron beam energy spread.

### S5. EXAMPLES OF CRYO-EM IMAGES OBTAINED WITH AND WITHOUT RELAY OPTICS

Commercially available apoferritin (VitroEase™ Apoferritin Standard, Catalog number: A51362, Thermo Fisher Scientific) proved to be a good specimen to use for comparing cryo-EM images recorded with and without relay optics (but both without a phase plate). As shown in Figure S2, particles are distributed quite uniformly, and in a desirably high abundance, when samples were used directly from the vial, as supplied.

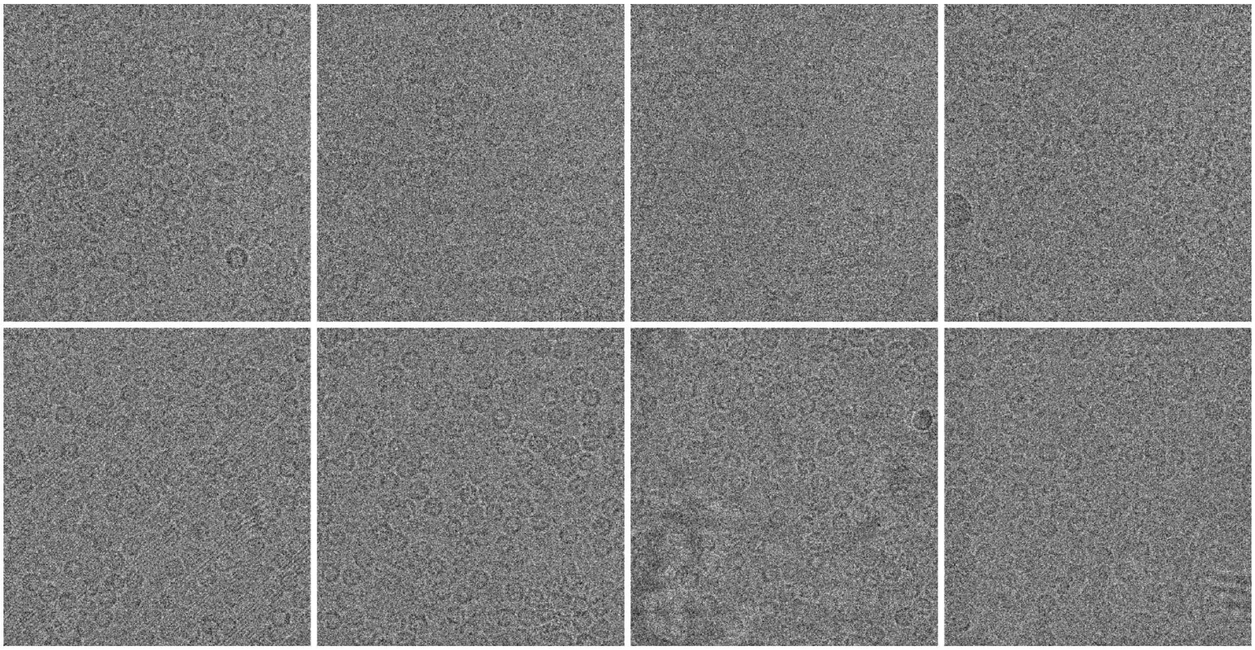

**Figures S2.** Representative cryo-EM micrographs of apoferritin obtained with (top row) and without (bottom row) relay optics. The width of the field of view is 179.7 nm with relay optics and 199.2 nm without relay optics. All images are displayed on a linear grayscale between 0.81 and 1.20, where 1.0 is the mean pixel value of each image.

### S6. JOHNSON NOISE CAN REMAIN A LIMITATION EVEN WHEN A MORE MONOCHROMATIC ELECTRON BEAM IS USED

Even when an 8 mm diameter electron beam hole is used in the laser phase plate (when using the relay optics), there is still a substantial amount of image blur from Johnson noise in the microscope's liner tube. Decreasing the electron beam energy spread would improve the CTF envelope, but due to the remaining, theoretically estimated Johnson noise, the resulting CTF envelope still falls short of what is found without relay optics. This is illustrated in Figure S3, which shows the theoretical CTF envelopes incorporating the effects of temporal coherence and Johnson noise both with and without relay optics for monochromated and non-monochromated electron sources. Methods for removing this limitation are discussed in Section 6.

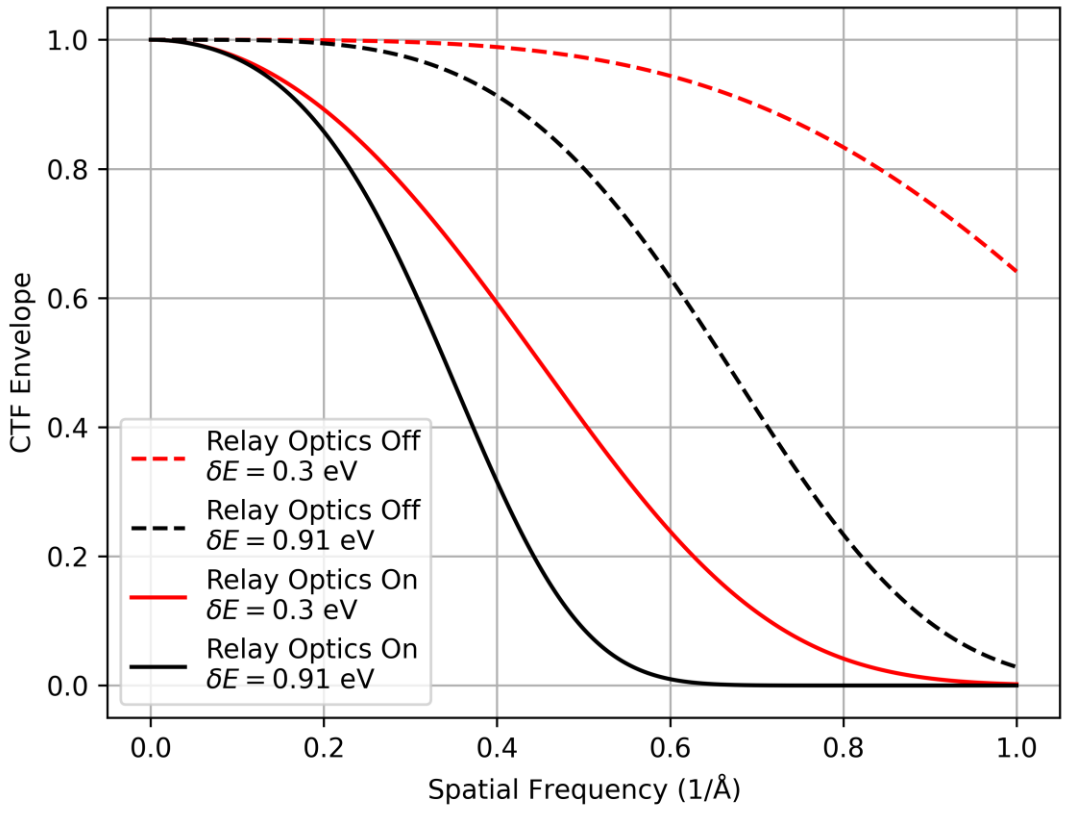


**Figure S3**. Expected improvement in our modified Titan's CTF envelope when using electrons with a narrower energy spread. The current CTF envelope with relay optics using the measured values of image blur variance $\langle \delta x^2 \rangle$ and electron beam energy spread $\delta E$ for the 8 mm electron beam hole diameter dummy (see Figure 3) is shown in the solid black line. The solid red line shows this case except that $\delta E$ is reduced to 0.3 eV. For comparison, the dashed black line shows the CTF envelope without relay optics using the measured value of $\delta E$, and the dashed red line shows the CTF envelope without relay optics if $\delta E = 0.3$ eV.
